# HEIP1 is required for efficient meiotic crossover implementation and is conserved from plants to humans

**DOI:** 10.1101/2022.12.22.521598

**Authors:** Dipesh Kumar Singh, Qichao Lian, Stephanie Durand, Aurelie Chambon, Aurelie Hurel, Birgit Walkemeier, Victor Solier, Rajeev Kumar, Raphaёl Mercier

## Abstract

Crossovers (CO) shuffle genetic information and physically connect homologous chromosome pairs, ensuring their balanced segregation during meiosis. COs arising from the major class I pathway require the activity of a well-conserved ZMMs group of proteins which, in conjunction with MLH1, facilitate the maturation of DNA recombination intermediates specifically into COs. The HEIP1 protein was identified in rice and proposed to be a new, plant-specific member of the ZMM group. Here we establish and decipher the function of the *Arabidopsis thaliana* HEIP1 homolog in meiotic crossover formation and report its wide conservation in eukaryotes. We show that the loss of Arabidopsis HEIP1 elicits a marked reduction in meiotic COs and their redistribution towards chromosome ends. Epistasis analysis showed that *AtHEIP1* acts specifically in the class I CO pathway. Further, we show that HEI1P acts both prior to crossover designation, as the number of MLH1 foci is reduced in *heip1*, and at the maturation step of MLH1-marked sites into COs. Despite the HEIP1 protein being predicted to be primarily unstructured and very divergent at the sequence level, we identified homologs of HEIP1 in an extensive range of eukaryotes, including mammals.

## Introduction

Accurate ploidy reduction during meiosis in sexually reproducing organisms relies on physical connections, known as chiasmata, between homologous chromosome pairs. Chiasmata are the cytological manifestation of genetic crossovers (COs) arising from homologous recombination during prophase I of meiosis. Failures or errors in CO formation are tightly linked with chromosomal missegregation leading to sterility or aneuploidy, such as Down syndrome in humans (1–3). COs, through reciprocal exchanges, also give rise to chromosomes with novel combinations of parental alleles, providing a major source of genetic variation.

During meiosis, CO formation is initiated by programmed induction of DNA double-strand breaks (DSBs)(4, 5). A minor fraction of these DSBs mature into COs through the action of two pathways (class I and class II) that coexist in many organisms (6–8). In most eukaryotes, including *Arabidopsis thaliana*, Class I COs constitute the major proportion of COs and are subject to CO interference, a poorly understood process that prevents the formation of two class I COs in close vicinity to each other (9, 10). The class I pathway depends on the activity of the conserved MLH1-MLH3 (MutLγ) nuclease that processes meiotic intermediates specifically into CO (11–13)(11–14). Cytologically, MLH1 forms foci between the homolog pairs in late prophase I that correspond to class I CO-designated sites (15, 16). Class II COs are dependent on the activity of structure-specific endonucleases, including MUS81, and can occur in close proximity to each other without interference (8, 17, 18).

The formation of class I COs is dependent on a group of remarkably well conserved meiosis-specific proteins collectively referred to as the ZMMs in *Saccharomyces cerevisiae*: <u>Z</u>ips (Zip1/Zip2/Zip3/Zip4), <u>M</u>sh4-Msh5, <u>M</u>er3, and Spo16. The ZMM proteins act as pro-crossover factors in different subcomplexes that recognize early DNA recombination intermediates and process progressively CO-specific intermediates during meiosis (19–21). The ZMM protein homologs in *A. thaliana* are ZYP1, SHOC1, HEI10, ZIP4, MSH4, MSH5, MER3, and PTD (22), and they are all, except for ZYP1, essential for class I CO formation, demonstrating the conservation of the pathway at the biochemical level (22). The exception, ZYP1, is a transverse element in the central region of the synaptonemal complex (SC), a tripartite structure that tethers homologous chromosomes during prophase I (23). Oppositely to other ZMMs, the Arabidopsis *zyp1* mutant displays an increase in the class I CO frequency and abolished interference, showing that ZYP1 is not required for CO formation but rather regulates their number and distribution (15, 24). The Arabidopsis and rice ZMM HEI10 are structurally and functionally related to the *S. cerevisiae* RING domain-containing Zip3 protein (25, 26). Both Arabidopsis and rice HEI10 proteins display a highly dynamic localization on meiotic chromosomes. HEI10 initially forms numerous discrete foci on synapsed chromosomes and progressively concentrates into a few prominent foci in late prophase I. These prominent HEI10 foci colocalize with the CO-designated site marked by MLH1 (15, 27). HEI10 is proposed to undergo a diffusion-mediated coarsening process leading to the formation of a few spaced out foci designating the CO sites (27, 28). In support of this model, HEI10 dosage regulates meiotic CO frequency in Arabidopsis, and overexpression of *HEI10 (HEI10-OE*) increases CO frequency over two-fold (29). Thus, HEI10 dynamics may directly regulate the CO positioning, although the detailed mechanism has yet to be described.

The rice HEI10 Interacting Protein 1 (HEIP1) was shown to interact in a yeast two-hybrid with HEI10, MSH5, and ZIP4 (30, 31). Rice *heip1* mutants exhibit a severe drop in chiasma frequency and a marked reduction in late HEI10 foci. Further, *heip1hei10* and *heip1* mutants show a similar reduction in chiasmata number. Altogether, this suggests that HEIP1 promotes class I COs. Rice HEIP1 colocalizes with HEI10 and displays the same dynamic localization on meiotic chromosomes: initially, numerous discrete foci are observed at early prophase I, followed by a few prominent foci at late prophase I. The localization of HEIP1P on the chromosome is dependent on HEI10 and ZIP4. The rice HEIP1 is thus proposed to be a novel ZMM protein, but earlier analysis failed to identify orthologs outside plants (30, 31).

Here, we established the role of the Arabidopsis HEIP1 (AtHEIP1) in meiotic CO formation. We analyzed a series of *Arabidopsis heip1* mutants, which exhibited defects in meiotic CO formation. Through epistatic interactions, we show that *AtHEIP* acts specifically in the class I CO pathway. Arabidopsis *Atheip1* mutants displayed a severe reduction in CO and bivalent formation but showed a rather small decrease in HEI10-MLH1 focus numbers. HEI10 overexpression yields an MLH1 foci formation in both wild type and *heip1* but did not increase CO frequency in the *heip1*. Altogether, this suggests that HEIP1 acts both upstream and downstream of the CO site designation process in the class I CO pathway. Furthermore, we identified HEIP1 homologs in a wide range of eukaryotes, including mammals, strongly suggesting that HEIP1 is a conserved pro-crossover factor with roles in CO maturation.

## RESULTS

### AtHEIP1 promotes chiasmata formation

The protein encoded by AT2G30480, which we here term AtHEIP1, is the sole *Arabidopsis* homolog of rice HEIP1 with 28% protein sequence identity between the full-length proteins (30, 31). We explored the function of AtHEIP1 by analyzing five *Atheip1* mutant lines, including three T-DNA insertion lines in Columbia-0 (Col-0 *heip1-1, heip1-2*, and *heip1-3*) and two full-deletion lines generated by CRISPR-Cas9 genome editing in the Col-0 (*heip1-4*) and Landsberg (Ler, *heip1-5*) ecotypes (Figure 1A & S1 dataset). The five *heip1* mutants displayed no visible growth or developmental defects. However, the fertility (seed per fruit) was substantially reduced compared to wild-type sister plants (Fisher’s LSD test, p<10^-6^, Figure 1B). We observed a similar level of fertility reduction in *heip1-1, heip1-2*, (two insertion lines) (p>0.5), and *heip1-4*, (a deletion line), suggesting that these three Col-0 lines are null alleles (Figure 1B). The *heip1-3* is slightly more fertile compared to the other three Col-0 lines (p<10^-4^) and is likely not a null allele. The fertility is slightly less affected in the Ler *heip1-5* deletion line compared to the Col-0 *heip1-4* deletion allele (Figure 1B), suggesting the HEIP1 could play a less prominent role in the Ler background.

**Figure 1.**
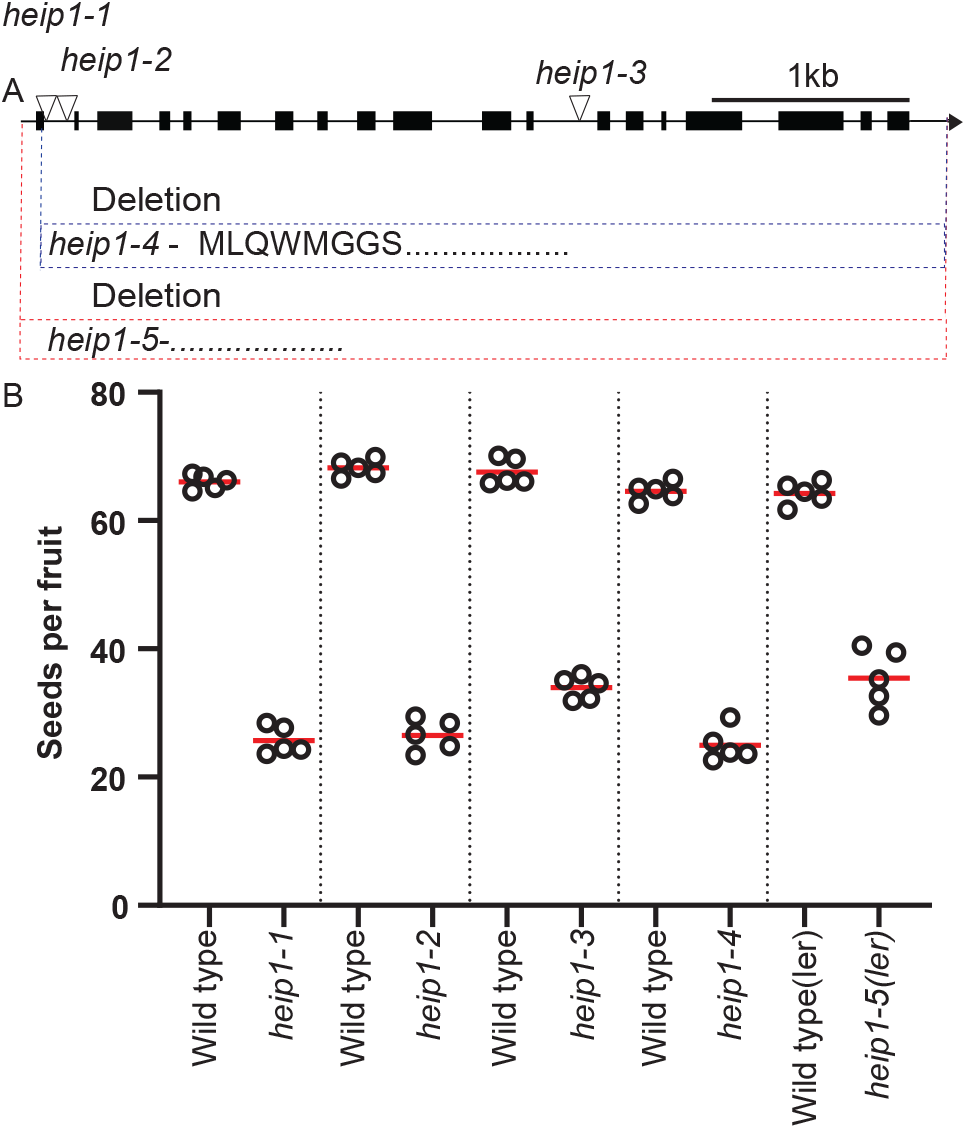
Schematic representation of the Arabidopsis *HEIP1* gene, *heip1* mutations, and fertility analysis of *heip1* mutants. A. The gene orientation is indicated by a horizontal arrow, and exons are indicated by solid black boxes, while introns and untranslated regions (UTR) are represented by the black line. Inverted triangles indicate T-DNA insertion points and black and red dotted lines denote the deleted region of *HEIP1* in Columbia.0 and Landberg’s ecotypes, respectively. The *heip1-4* allele may synthesize only eight amino acids represented by a single letter code, whereas the *heip1-5* has a complete deletion of the coding sequence. B. Each black circle represents the average seeds per fruit for one plant, obtained by counting at least ten fruits per plant. Comparison of fertility based on the number of seeds per fruit in a series of *heip1* mutants. The mean for each genotype is represented by a red bar. Each *heip1* mutant allele is compared with wild-type sister plants that were cultivated together in a segregating population.

We performed chromosome spreads to determine the role of AtHEIP1 during meiosis. Spreads of male meiocytes revealed that meiotic progression in *heip1* mutants is similar to wild type until the pachytene stage, where chromosomes appear fully synapsed (compare Figure 2A to 2G). At diakinesis, chromosome condensation revealed the presence of five bivalents in the wild type, with homologous chromosomes connected by chiasmata (Figure 2B). In contrast, a mixture of bivalents and univalents were observed at the diakinesis stage in *heip1* mutants (Figure 2H), suggesting a defect in CO formation. At metaphase I, five bivalents align on the metaphase plate in wild-type cells (Figure 2C). An average of 2.5 univalent pairs and 2.5 bivalents per cell were observed in *heip1-1* (Figure 2I), with similar numbers obtained in *heip1-2, heip1-4 and heip1-5* compared to *heip1-1*, (Uncorrected Dunn’s test p=0.34, 0.65, 0.28 respectively; Figure 2M). The *heip1-3* allele appeared less affected, with 3.8 bivalents per meiotic cell (p<10^-6^), corroborating the milder fertility defect of this line. As a probable direct consequence of the presence of univalents, unbalanced chromosome segregation and unequal nuclei were observed in subsequent meiotic stages (Figure. 2K-L). Female meiocyte spreads also revealed the presence of univalents in *heip1-1* (Mean = 2.9 bivalents per cell, Figure S1). The presence of univalents indicates that *Arabidopsis* HEIP1 is essential for the formation of wild-type levels of COs during male and female meiosis.

**Figure 2.**
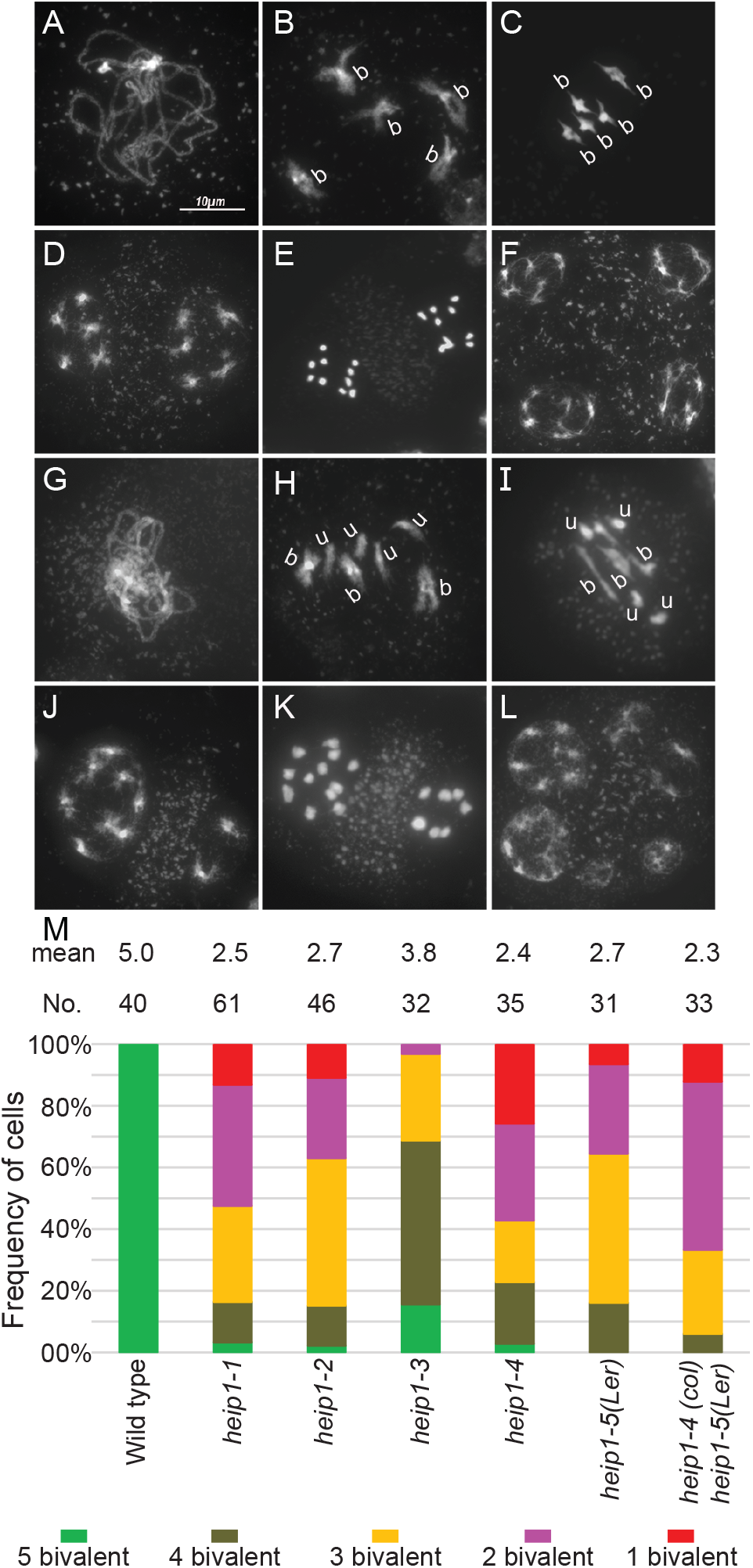
Chromosomes spreads reveal meiotic defects in *heip1* mutants. Wild type (A–F): (A) Pachytene. (B) Diakinesis and (C) metaphase I with five bivalents (b). (D) telophase I, with two sets of five chromosomes. (E) anaphase II. (F) tetrad. *heip1-1* (G–L): (G) Pachytene, which is indistinguishable from wild type. (H) Diakinesis with three bivalents (b) and two pairs of univalents (u). (I) Metaphase I with three bivalents and two pairs of univalents, (J) Telophase I with unbalanced chromosome distribution. (K) Anaphase II with unequal chromosome segregation, and (L) polyads. (Scale bar, 10 μm.). (M) Quantification of bivalents and univalents at metaphase I. Cells were categorized according to the number of pairs of univalents/bivalents. The mean bivalents number per cell and the number of cells analyzed are indicated above each bar.

### AtHEIP1 acts specifically in the class I CO pathway

Since AtHEIP1 appears to be involved in CO formation, we explored its epistatic relationship with key factors of class I and class II CO pathways. MSH5 and HEI10 are two canonical ZMM proteins required for class I CO formation. The Arabidopsis *hei10* and *msh5* mutants display a strong reduction of bivalents (1.5 mean bivalent per cell, Figure 3), more severe than *heip1-1* (mean= 2.5, p=0.011 and p=0.0003). Combining the *heip1-1* mutation with either *msh5* or *hei10* did not further reduce the bivalent numbers compared to single *msh5* and *hei10* (p=0.78 and p=0.96, Figure 3A, 3B&G), suggesting that HEIP1 acts in the same pathway as MSH5 and HEI10, though having a less important role.

**Figure 3.**
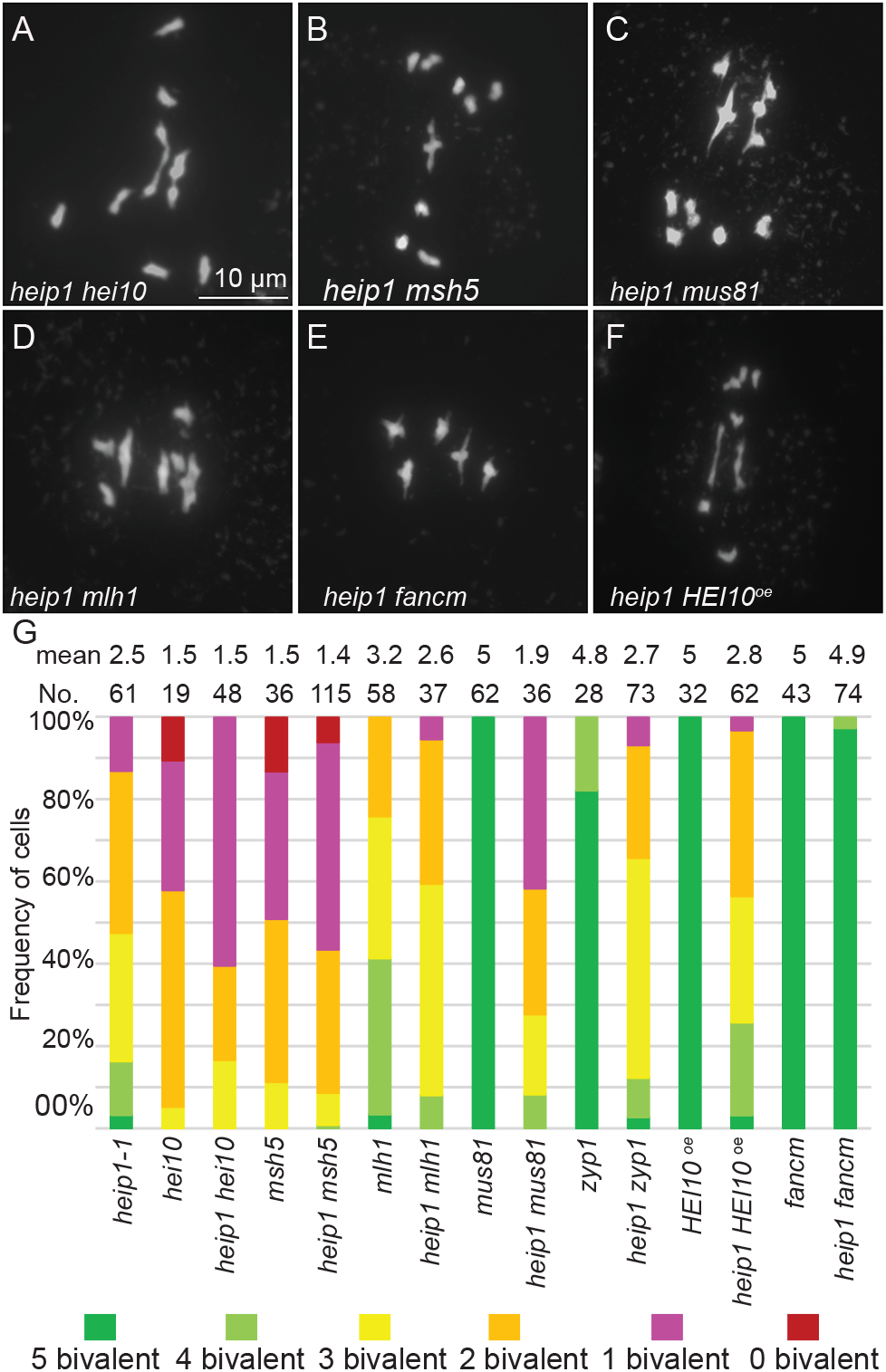
Epistasis analysis of *heip1-1*. (A–F) Representative image of metaphase I chromosome spreads of male meiocytes in the following genotypes: (A) *heip1-1 hei10*, (B*) heip1 msh5* (C) *heip1-1 mus81* (D) *heip1-1 mlh1* (E) *heip1-1 fancm* (F) *heip1-1 HEI10-OE* Scale bar = 10μm. (G) Quantification of bivalents at metaphase I. Cells were categorized according to the number of bivalents. The average number of bivalents per cell and the number of analyzed cells are indicated above each bar.

MLH1 is also involved in the class I CO pathway, but *mlh1* has a milder CO defect than *msh5* or *hei10* (Figure 3, p<10^-6^ and p=0.02, respectively), with an average of 3.2 bivalents, presumably because MLH1 acts later in the pathway. The *heip1* mutant is slightly more affected in bivalent formation than *mlh1* (p<10^-6^), and the double *heip1 mlh1* is not further reduced compared to the single *heip1* (p=0.69), suggesting that HEIP1 and MLH1 act in the same pathway, with HEIP1 acting upstream of MLH1 (Figure 3D&G). It was previously shown that overexpression of HEI10 in the wild type doubles the numbers of class I COs (29). However, HEI10 overexpression (*HEI10-OE*) in *heip1* did not significantly increase bivalent formation per cell (2.8 vs. 2.5 p=0.3), further suggesting that the class I pathway is defective in *heip1* (Figure 3F&G). Altogether, our genetic analysis suggests that AtHEIP1 acts in the same pathway as HEI10, MSH5, and MLH1 for class I CO formation.

To determine whether AtHEIP1functions in the class II CO pathway we mutated the MUS81 endonuclease, which generates class II COs (17, 32), in the *heip1-1* background. The *heip1-1 mus81* double mutant showed a lower bivalent count compared with *heip1-1* mutants (Figure 3C&G; bivalents, 1.9 vs. 2.5, p=0.047), supporting the conclusion that HEIP1 and MUS81 act in different pathways to promote CO formation. To further test whether the class II pathway is functional in *heip1-1*, we combined *heip1-1* with *fancm*, a mutation causing a massive increase in class II COs (33). The introduction of the *fancm* mutation in *heip1* provoked an almost complete restoration of mean bivalent formation (Figure 3E and 3G), arguing that the class II pathway is functional in the absence of HEIP1. Thus, HEIP1 appears to be dispensable for the class II CO pathway and is specific to the class I pathway.

### AtHEIP1 is dispensable for synapsis but required for normal formation of HEI10/MLH1 foci

We further investigated the role of Arabidopsis HEIP1 in synapsis and CO formation by performing immunostaining of the axial element (REC8), the transverse element of the SC (ZYP1), and class I CO factors (HEI10 and MLH1) (Figure 4 & 5). In the wild type, REC8 forms a continuous signal along the chromosome axes in the zygotene, pachytene, and diplotene stages (Figure 4 I–L). Synapsis initiates at zygotene, with stretches of ZYP1 zipping the two homologous axes together (Figure 4A) and extending to the entire chromosome length at pachytene (4B–C). Multiple HEI10 foci decorate the synapsed regions at zygotene (Figure 4E) and early pachytene (Figure 4F). During pachytene, the HEI10 signal along the synaptonemal complex is progressively less homogenous, with the coarsening of HEI10 foci (Figure 4G) (27, 28). At diplotene, chromosome desynapse with a complete disappearance of the ZYP1 signal, while HEI10 appears as large foci, marking class I CO-designated sites (Figure 4 D, H) (26, 28). In *heip1-1*, the localization of REC8, ZYP1, HEI10, and MLH1 was qualitatively indistinguishable from the wild type (Figure. 4M–X), showing that Arabidopsis HEIP1 is dispensable for synapsis and HEI10 loading onto the synapsed chromosomes. The coarsening of HEI10 foci appears also to be unaffected in *heip1* (Figure 4S), with large HEI10 and MLH1 foci colocalizing at late prophase I (Figure 5). The quantification of the HEI10/MLH1 co-foci at the diplotene/diakinesis stage (Figure 5U) revealed a slight reduction in *heip1-1* compared to wild type (from 11.6 to 9, Fisher’s LSD test p<0.0001), *heip1-4* (from 12.2 to 10.2, p=0.0017) and the Col/Ler hybrid *heip1-4/heip1-5* (From 9.4 to 7.1, p<0.0001). These data show that HEIP1 is not essential for synapsis and the recruitment of HEI10 to synapsed chromosomes but is needed for the formation of a normal number of HEI10/MLH1 cofoci.

**Figure 4.**
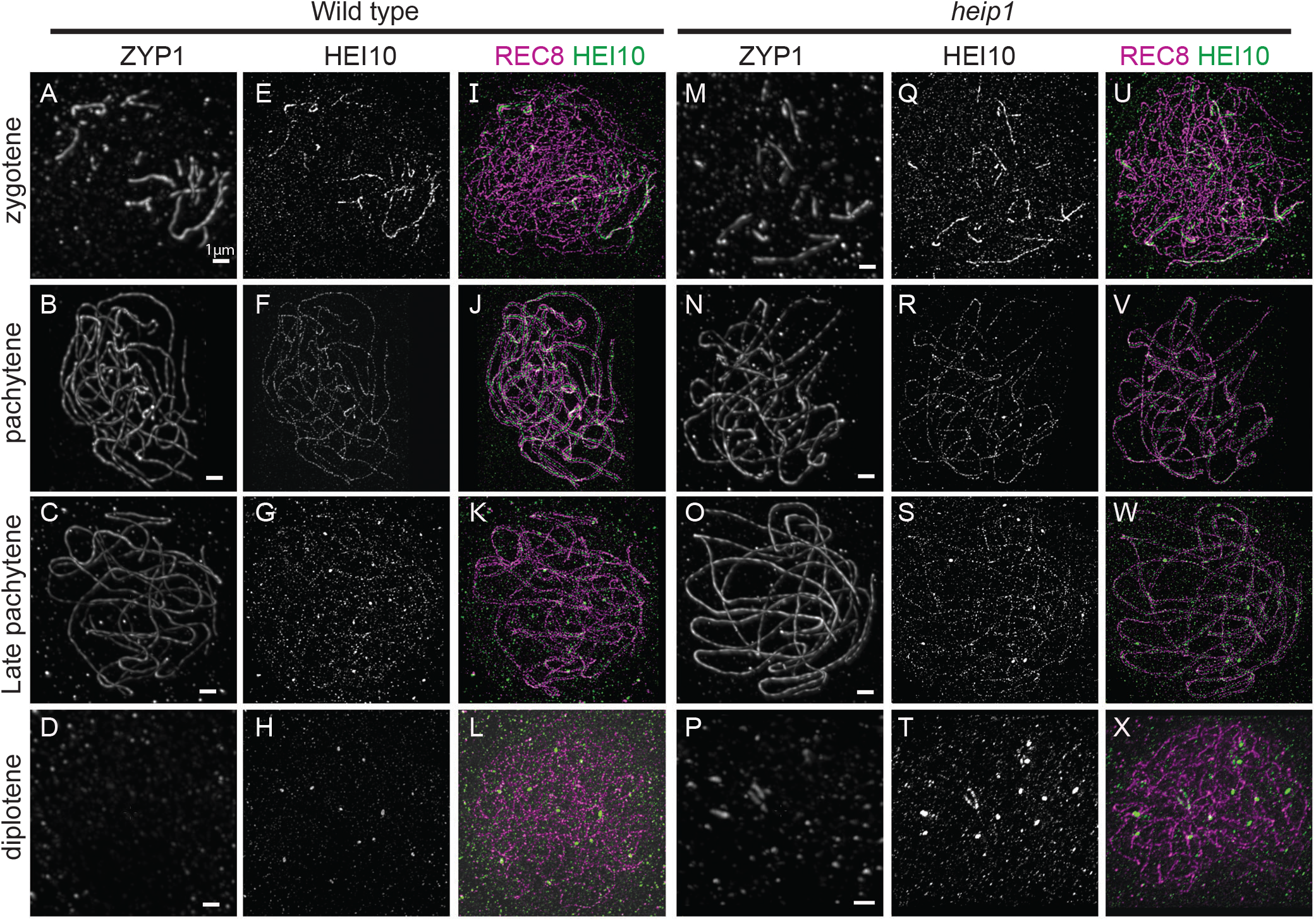
Synapsis and HEI10 dynamics are unaffected in *heip1-1*. The three columns on the left panel represent wild-type meiocytes and the three columns on the right panel show *heip1-1* meiocytes in various stages of meiotic prophase I at zygotene (A, E, I, M, Q, & U), pachytene (B, F, J, N, R, & V), late pachytene (C, G, K, O, S, & W), and diplotene (D, H, L, P, T, & X). Immunolocalization of REC8 (purple, E-H & Q-T), HEI10 (green, I-L & U-X) and ZYP1 (white, A-D & M-P) on male meiocytes. ZYP1 was acquired with confocal microscopy, while HEI10 and and REC were acquired with STED microcopy. Scale bar =1μm.

**Figure 5.**
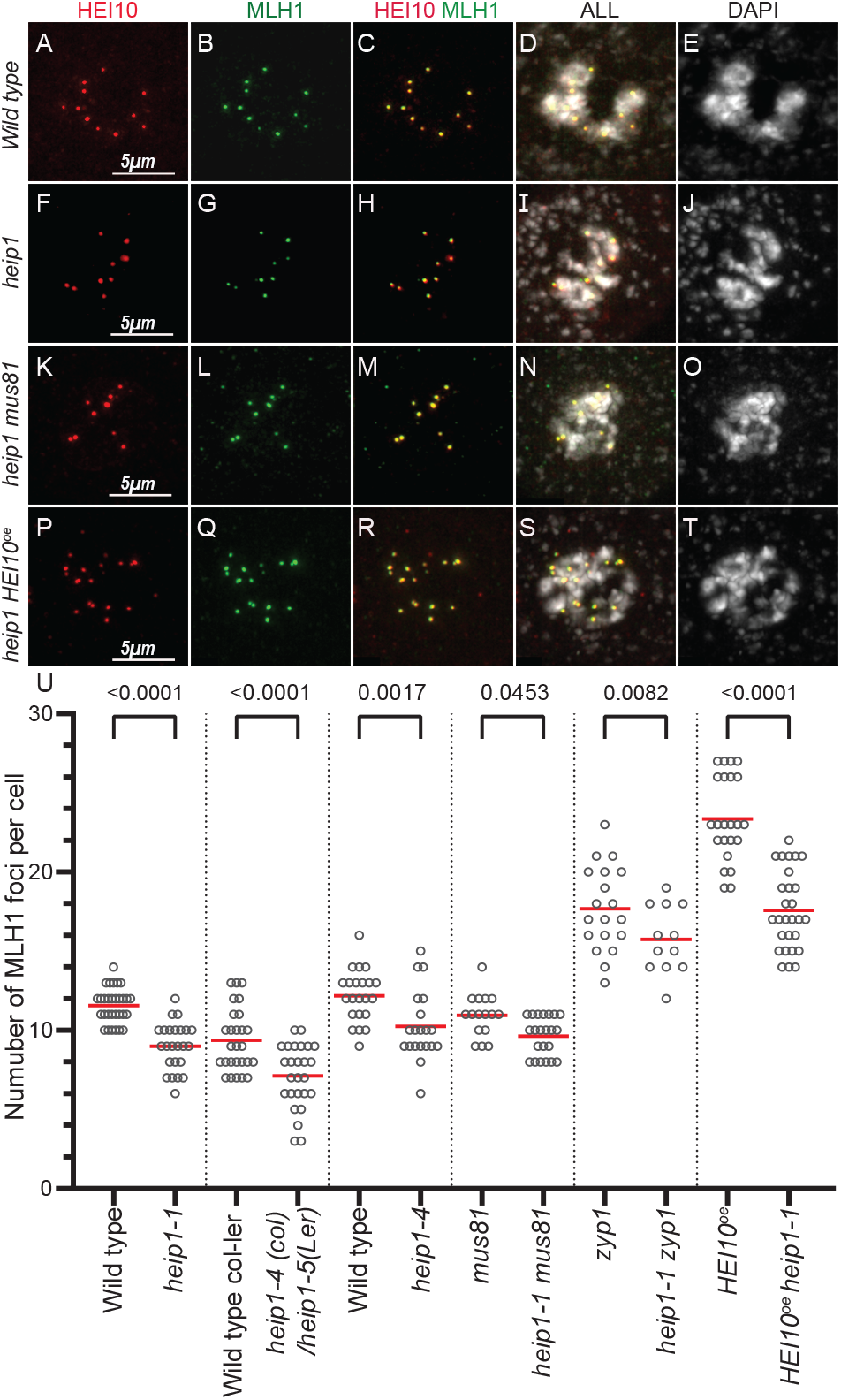
The number of HEI10-MLH1 foci is reduced in *hei1p*. Immunolocalization of MLH1 and HEI10 on male meiocytes at diplotene/diakinesis. (A-E) Wild type. (F-J) *heip1*. (K-O) *heip1 mus81*. (P-T) *heip1 HEI10-OE*. From left to right: HEI10 in red (A, F, K, and P), MLH1 in green (B, G, L, and Q), merge HEI10 & MLH1 (C, H, L, and Q), merge All (D, I, N, and S), and DAPI in white (E, J, O, and T). (U) Quantifications of MLH1–HEI10 cofoci. Each mutant was compared to sibling controls and separated by vertical lines. Each dot is an individual cell, and the red bar is the mean. Scale bar = 5 μm. p values are from Fisher’s LSD tests.

### HEIP1 acts both upstream and downstream of MLH1 focus formation

In the wild-type (Col-0) male meiosis, an average of 11 MLH1 foci per cell are converted into chiasmata that provide physical connections between homolog pairs, resulting in five bivalents per cell at metaphase I. In *heip1*, we noticed a discrepancy between the mean MLH1 foci number during prophase I and bivalent formation at metaphase I: The reduction of HEI10/MLH1 foci in *heip1* mutants appears modest (22%) compared to the observed frequency of univalents (50% of the chromosomes lacking COs). A formal possibility is that HEI10/MLH1 co-foci and COs cluster on some homolog pairs, while other chromosomes lack foci and COs. However, tracing chromosomes at late pachytene in *heip1* did not indicate the clustering of HEI10/MLH1 co-foci and showed that each chromosome receives at least one HEI10/MLH1 co-focus (Figure 6, n=5 cells). The numerous univalents together with the observation that all chromosomes are decorated by at least one focus strongly suggests that a large proportion of HEI10/MLH1 foci fail to mature into COs in absence of HEIP1. Two additional observations further support this interpretation: (i) The number of HEI10/MLH1 foci is similar in *heip1 mus81* and *heip1* (Figure 5, p=0.27; 9.6 and 9 per cell, respectively), but numbers of bivalent are reduced in *heip1 mus81* (average 1.9 per cell) compared to *heip1* (average 2.5 per cell, Figure 3). This larger excess of HEI10/MLH1 foci relative to bivalent numbers in *heip1 mus81* further suggests a defect in conversion of MLH1 foci into COs. (ii) HEI10 overexpression (*HEI10-OE*) or *zyp1* mutation both increase the numbers of HEI10/MLH1 foci and class I COs (15, 24, 29) (Figure 5&S2). In the *heip1* background, HEI10 overexpression and *zyp1* mutation also elicited a massive increase in MLH1 foci, which reached an average of 17.6 in *heip1-1 HEI10-OE* and 15.6 in *heip1-1 zyp1*. This is lower than in the corresponding single *HEI10-OE* (23.3, p<10^-6^) or *zyp1* (17.7, p=0.008), further showing that HEIP1 contributes to athe normal level of HEI10/MLH1 foci formation. In sharp contrast, the number of bivalents was not significantly increased in *heip1-1 HEI10-OE* (average 2.8 per cell) and *heip1-1 zyp1* (2.7 per cell) compared to *heip1-1* (2.5 per cell, p=0.31 and p=0.36), resulting in a marked discrepancy between the HEI10/MLH1 foci number and the number of bivalents. Altogether, this shows that HEI1P plays a crucial role in converting MLH1 foci into COs, in addition to its upstream role in promoting HEI10/MLH1 focus formation.

**Figure 6.**
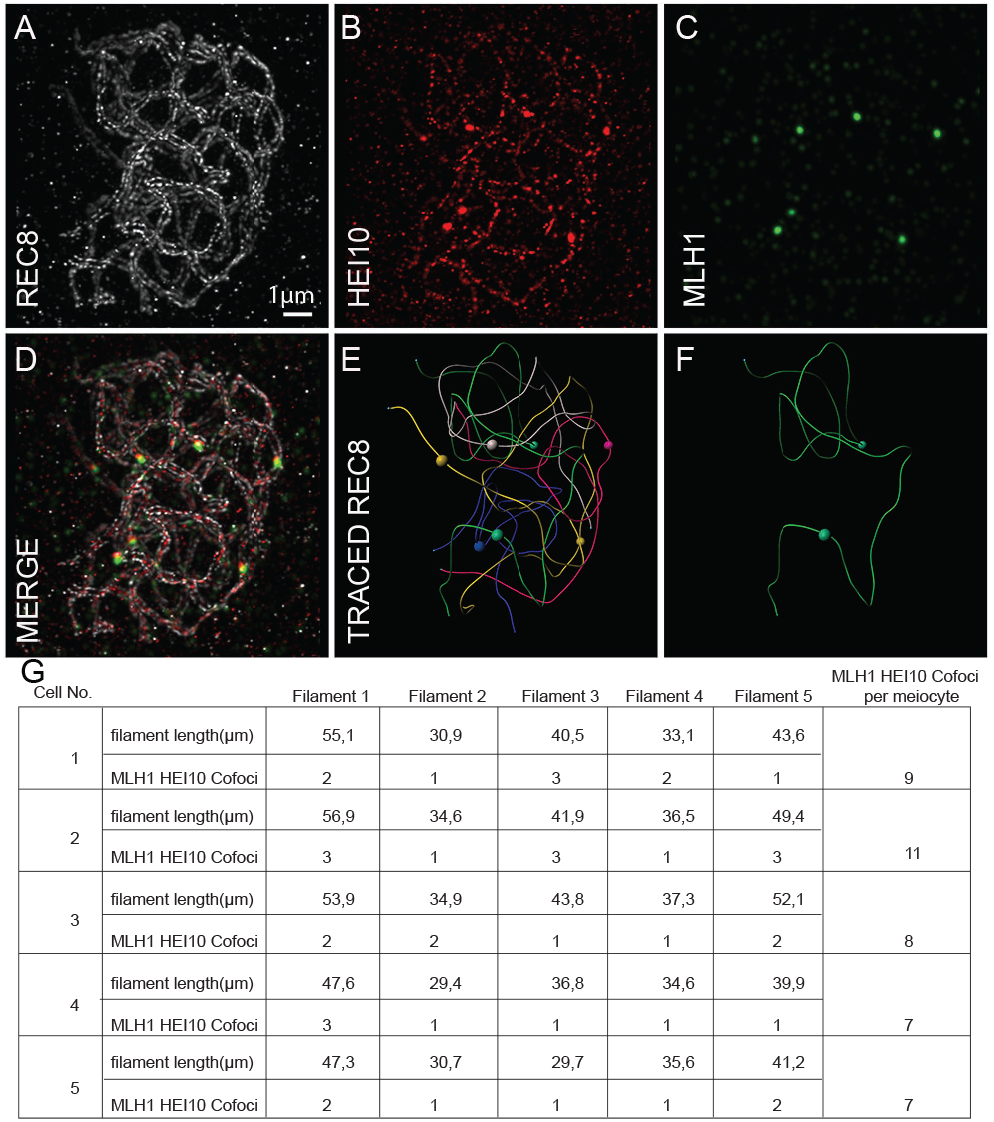
Analysis of the distribution of HEI10-MLH1 foci in *heip1*. (A–D) Triple immunolocalization of REC8, HEI10, and MLH1 on *heip1* male meiocytes. Imaging was done with 3D-STED and the projection is shown. Scale bar=1 μm. (E and F) REC8 signal was traced in 3D using the IMARIS tool. (E) All five bivalents are represented in a separate color and a similar color was used to mark HEI10/MLH1 co-foci on that bivalent. (F) Representation of one bivalent with HEI10/MLH1 co-foci. (G) The length of chromosomes (filaments) and distribution of HEI10/MLH1 foci among chromosomes analyzed in five cells are presented in a table.

### Aneuploids are common among heip1 progeny

To measure COs genetically, we produced F1 hybrids carrying two deletion *HEIP1* alleles, *heip1-4* (Col-0 strain) and *heip1-5* (Ler strain). These F1s, and sibling wild-type controls, were crossed as male or female to wild-type Col-0 and the progeny were whole-genome-sequenced with short reads. Analyses based on sequence coverage and allelic ratio (Supplementary dataset 2) detected 10.1% and 18.7% of trisomics in progeny derived from *heip1* female and male, respectively, while none were detected in a total of 817 wild-type progeny (Table 1). The two shortest chromosomes, 2 and 4, were overrepresented (41% and 38% of the trisomy), while trisomy 3 and 5 were rarer (8% and 13%), and trisomy 1 was absent. In addition, some double-trisomics (Trisomy 2+4 and trisomy 2+5) and triploids were detected (Table 1). The trisomy is likely derived from the unbalanced segregation of achiasmatic chromosomes, as supported by the absence of COs on the trisomic chromosome in all 78 trisomic samples, and the observation that the trisomic genotype is systematically Ler/Col-0/Col-0 (Ler/Col-0 from the *heip1* gamete, Col-0 from the Col-0 parent), supporting a failure to distribute homologs at meiosis I rather than chromatids at meiosis II. The different relative contributions of the five chromosomes to aneuploidies could be due to both crossover frequency (i.e., small chromosomes are more prone to lack COs) and/or different transmission rates of the aneuploid gametes. Furthermore, ten cases of genomic rearrangements were detected in the progeny of *heip1* females, with the addition of half of a chromosome, and/or a more complex pattern (Supplemental figure 2) that could correspond to chromoanagenesis(34).

**Table 1:**
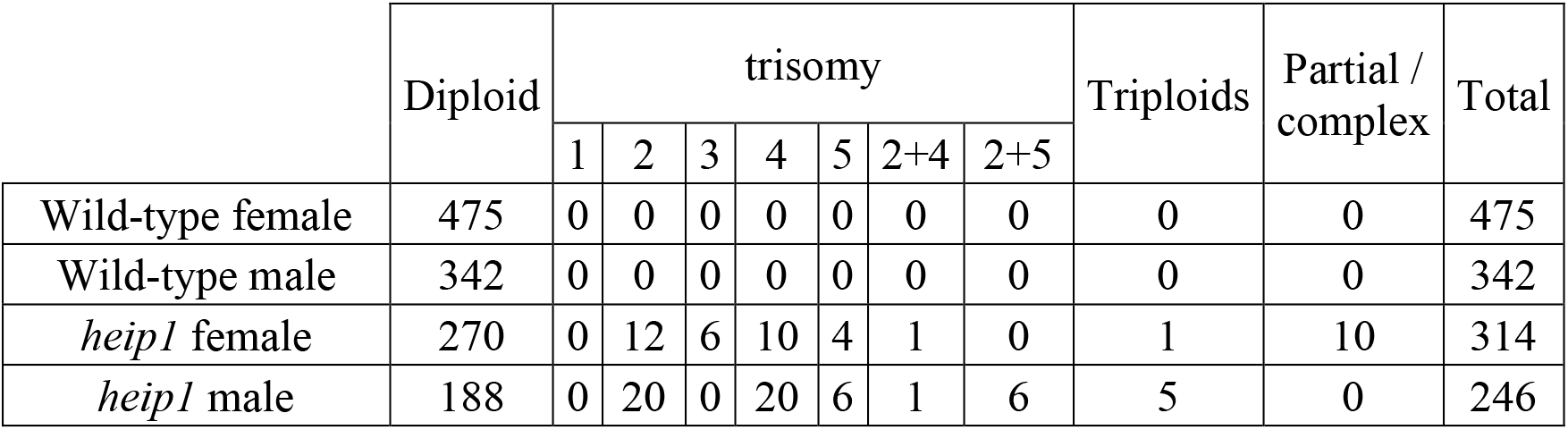
Analysis of aneuploidy in wild type and *heip1* mutant offspring. The numbers of trisomies, triploids and other complex events are presented.

|  | Diploid | trisomy |  |  |  |  |  |  | Triploids | Partial / complex | Total |
| --- | --- | --- | --- | --- | --- | --- | --- | --- | --- | --- | --- |
|  |  | 1 | 2 | 3 | 4 | 5 | 2+4 | 2+5 |  |  |  |
| Wild-type female | 475 | 0 | 0 | 0 | 0 | 0 | 0 | 0 | 0 | 0 | 475 |
| Wild-type male | 342 | 0 | 0 | 0 | 0 | 0 | 0 | 0 | 0 | 0 | 342 |
| *heip1* female | 270 | 0 | 12 | 6 | 10 | 4 | 1 | 0 | 1 | 10 | 314 |
| *heip1* male | 188 | 0 | 20 | 0 | 20 | 6 | 1 | 6 | 5 | 0 | 246 |

### Genetic crossovers are reduced and shifted to chromosome ends in heip1

Progeny sequencing revealed that the average number of genetic COs per transmitted male gamete was drastically reduced from 5.38±1.94 (mean ± SD) in wild type to 1.08 ± 1.29 in *heip1* (Figure 7A), confirming the crucial role of HEIP1 in crossover formation. Intriguingly, the number of observed genetic crossovers was even lower than expected according to the observed number of bivalents in *heip1*. In females, the number of COs was reduced from 2.8±1.29 to 2.46±1.37 (p=0.0013). The apparent reduction of crossover number in females is modest and may be due to the counter-selection of achiasmatic chromosomes, but the distribution of crossovers along chromosomes is markedly different compared to wild type (Figure 7B). Notably, the proximal regions which are the highest recombining regions in wild-type females are “colder” in *heip1*, and conversely, the terminal regions that have a low frequency of COs in wild type, exhibit increased numbers of COs in *hei10* females (Figure 7B). The low numbers of COs observed in male *hei10* prevents meaningful analysis of crossover distribution along chromosomes. We next explored CO interference in females, using coefficient of coincidence curves (Figure 7C). CO interference is still detected but appears to be slightly reduced in female *heip1* compared to wild type. The presence of interference among COs in *heip1* is consistent with a late role of HEI1P, downstream of CO site designation (35).

**Figure 7.**
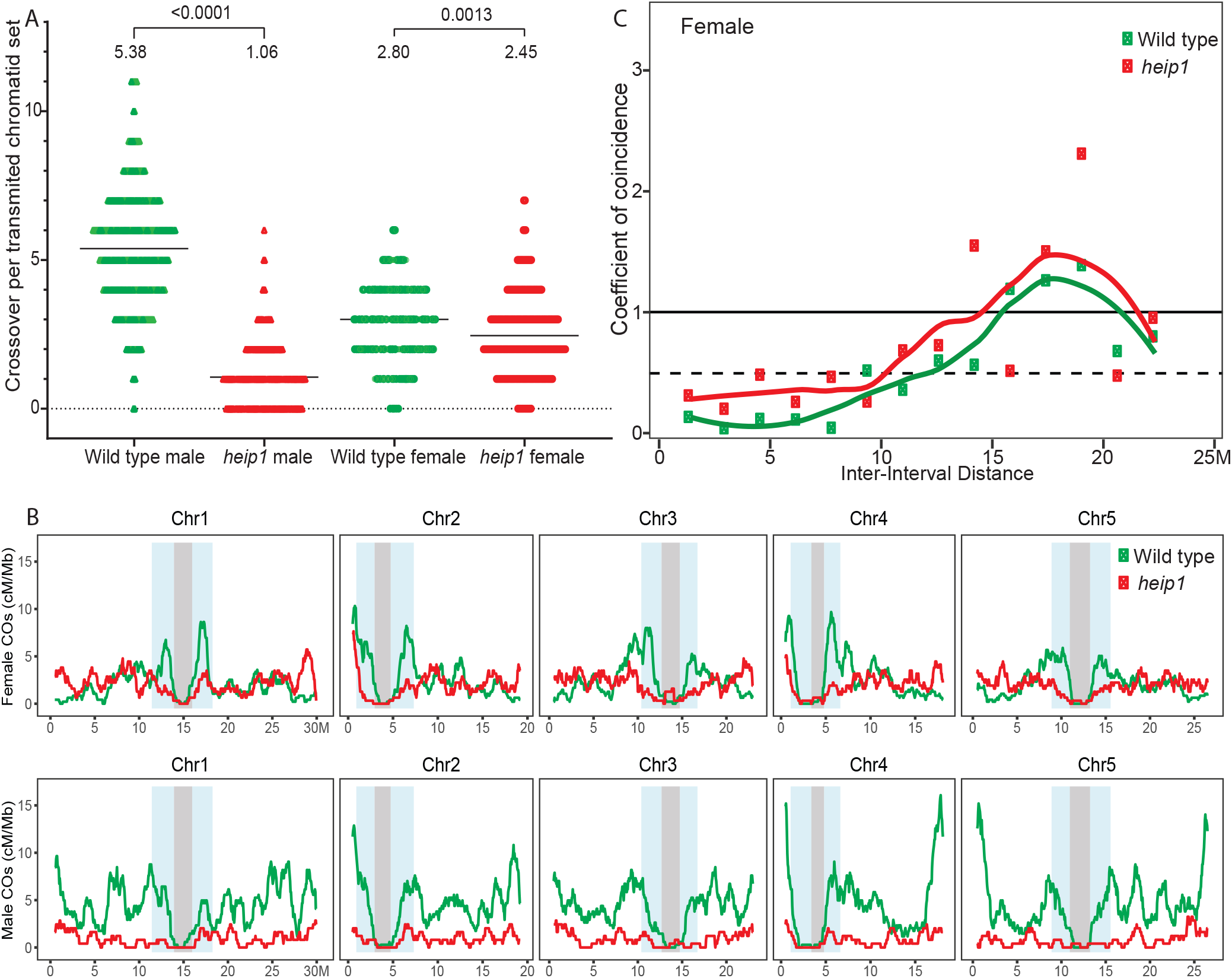
Analysis of CO numbers, distribution, and interference. (A) The number of COs per transmitted chromatid in female and male gametes of wild type and *heip1*. Fisher’s LSD test was applied (B) The chromosomal distribution of COs in female and male wild type and *heip1*, with a window size of 1 Mb and step size of 50 kb. (C) CoC curves in female and male meiosis of wild type and *heip1*, in which chromosomes were divided into 15 intervals to estimate the mean coefficient of coincidence.

### HEIP1 appears to be conserved in a large range of eukaryotes

Searching for homologs of HEIP1 in other species using classical tools such as BLAST retrieved exclusively plant proteins, the most distantly reported homologs of rice and Arabidopsis HEIP1 being found in nonvascular plants such as mosses, liverworts, and algae (31). Here, using HHblits with sensitive parameters (E-value cutoff for inclusion 0.02; Max target hits 1000; Number of iterations 7, Min probability on hitlist 20%) (36, 37). and Arabidopsis HEIP1 (Uniprot F4INT5) as a bait, we retrieved typically one hit per species in an extensive range of eukaryotes, including the human protein C12orf40 and its homolog in other vertebrates. C12orf40 is very strongly expressed in testis, supporting a potential meiotic role (EMBL-EBI expression Atlas). Conversely, using C12orf40 as bait we retrieved a single hit in a large range of animals and retrieved the HEIP1 plant homologs. HHblits detected sequence similarity between plants and animals only in the first ~70 N-terminal amino acids of the proteins (Figure 8 and sup dataset 3), a short region that is predicted to be structured by Alphafold, while the rest of the protein is predicted to be unstructured (38, 39). Using the Arabidopsis HEIP1 or the first 70 amino acids of human C12orf40 as bait in HHblits, we retrieved largely overlapping plant and animal proteins for diverse species with high scores (e.g., Arabidopsis HEI1P retrieved Human C12orf40 with E-value=4.2e-19) and additional hits in some fungal species (supplementary dataset 3). Within the conserved sequence patch, some amino acids are fully conserved between plants and animals (Figure 8), suggesting that they might play a key role in HEIP1 function. The orthology between plant HEIP1 and mammalian C12orf40 is further supported by the recent description of the crucial role of C12orf40 in promoting meiotic crossover formation in humans and mice (40). Altogether, this strongly suggests that HEIP1 and its meiotic function are largely conserved in eukaryotes.

**Figure 8.**
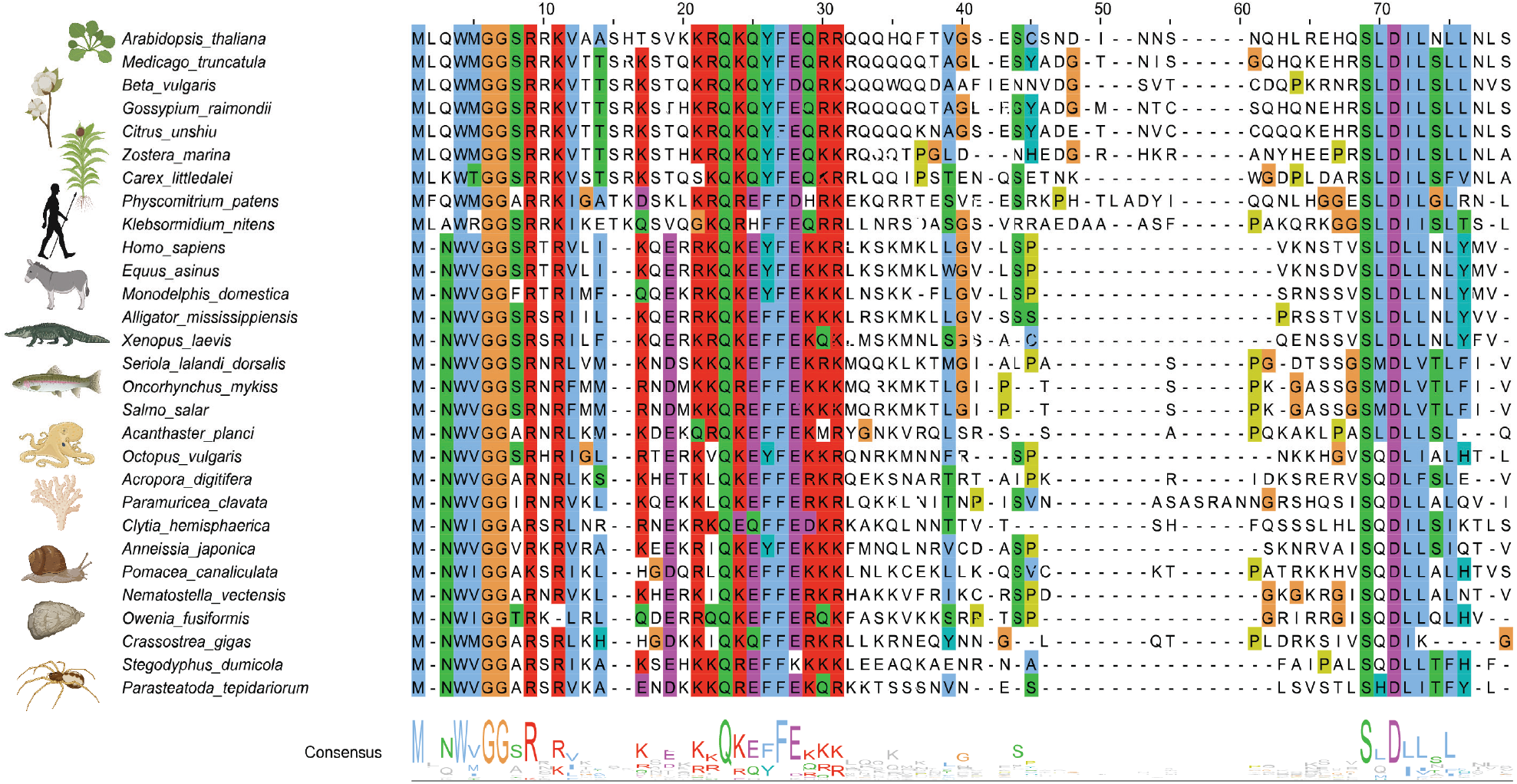
Sequence similarity between Arabidopsis HEIP1 and Human C12orf40 together with other plant and animal homologs. Homologs of HEIP1 and C12orf40 were retrieved with HHlits (Supplementary dataset 3). The N-terminal ends of proteins from representative species of diverse plant and animal clades were aligned in Jalview 2.11.2.5 using T-coffee with default parameters. Figure prepared with Biorender.

## Discussion

### Arabidopsis HEIP1 is a pro-class I crossover factor

Here, we have described the role of the Arabidopsis HEIP1 homolog in meiotic CO formation and have shown that AtHEIP1 is required for a wild-type level of chiasmata and COs. AtHEIP1 acts exclusively in the class I pathway together with ZMMs and is not involved in the formation of class II COs. The canonical Arabidopsis ZMM mutants *msh4, msh5, hei10, shoc1*, and *ptd1* have ~15% of wild type chiasmata levels, with the *mer3* mutant being slightly less affected with ~25% of wild-type levels (22, 41). In the same pathway, COs are reduced by ~40% in *mlh1/3* mutants, presumably as these proteins act later in the class I pathway (12, 22). The full deletion *Arabidopsis heip1-4* allele and two additional likely null alleles, display a roughly 50% reduction in chiasmata, which indicates a weaker CO defect compared to other *zmm* mutants. Moreover, *Arabidopsis heip1 hei10* and *heip1 msh5* mutants exhibited a lower number of chiasmata compared to *heip1*, which was at the level of *hei10* and *msh5*, indicating that the class I CO pathway is still partially operational in *heip1*. This suggests that HEIP1 is a ZMM promoting class I COs in *Arabidopsis*, but with a less critical role than the canonical ZMMs.

### Evolutionary conservation of HEIP1 in eukaryotes

Previous studies suggested that HEIP1 is a plant-specific single-copy gene family (30, 31). HEIP1 homologs could be straightforwardly detected in three clades of monocots, eudicots, and bryophytes (basal plants). Protein sequence analysis of plant HEIP1 homologs identified a highly conserved N-terminal region of ~250 amino acids with four conserved motifs of unknown function (31). Here, we this identified HEIP1 homologs outside plant species (Figure 8 and Supplementary dataset 3). The significant similarity between plant and animal homologs is restricted to the first ~75 N-terminal amino-acids, with some amino-acids having of 100% conservation (Figure 8), suggesting the existence of a crucial interaction interface. AlphaFold-based modelling of HEIP1 proteinspredicts a largely disordered or unstructured protein with only small patches forming helices. Disordered regions are relatively accessible and can potentially bind to multiple partners (42), which would be consistent with the reported interaction of HEIP1 with HEI10, MSH5, and ZIP4 (30, 31, 43). It is thus reasonable to speculate that HEIP1 could play a structural rather than an enzymatic role in implementing COs.We identified the *C12orf40* gene as the homolog of HEIP1 in humans, whose high expression in testis is compatible with a meiotic role. During the course of this study, a homozygous frameshift mutation in the *C12orf40* gene was identified in infertile men showing meiotic arrest (40). Mouse models mimicking the mutations in *C12orf40* from infertile patients show arrested spermatogenesis with a reduction in meiotic COs and bivalent formation (40). The reduction of MSH4 and TEX11/ZIP4 foci in the mutant spermatocytes points towards a defective class I CO pathway. Taken together, HEIP1 appears to be an evolutionarily conserved proteinpromoting the formation of class I COs that was likely present in the common ancestors of all living eukaryotes together with the ZMMS.

### Dual role of HEIP1 in CO formation

The main pathway of meiotic CO formation is largely conserved across the eukaryotic tree of life (4, 22, 44). In this pathway, the repair of double-strand breaks *via* inter-homolog interactions generates DNA precursors for CO formation. A subset of these precursors is designated before maturing into COs. The designation process ensures at least one obligate CO per chromosome and enforces CO interference, ensuring that COs are distributed away from another on the same chromosome. HEI10 localization is remarkably dynamic during prophase, with initial multiple foci along chromosomes subsequently consolidating into a small number of large foci that mark CO-designated sites (27, 28). HEI10 is thus considered as a marker of the CO designation process. Recently, HEI10 was proposed to play a more active role in the process and to drive CO designation through diffusion-mediated coarsening (27, 28). In late prophase, large HEI10 foci are decorated with MLH1, which mediates the final step of CO maturation with each MLH1 focus generating a CO (15, 26). While mechanistic details of CO maturation are still unclear, inefficiency in maturing MLH1-marked sites into COs is a major source of chromosome mis-segregation and aneuploidy in human female oocytes (35). Our data implicates AtHEIP1 both in the designation of CO sites and in the maturation of CO-designated sites into COs: (i) In the absence of AtHEIP1, the HEI10 dynamic appears qualitatively unaffected but results in a reduction in the HEI10/MLH1 co-foci numbers by ~20%. This indicates that AtHEIP1 plays an important, albeit non-essential, role in ensuring that the appropriate numbers of HEI10/MLH1 foci are formed during meiosis. (ii) We detected a CO maturation failure with a discrepancy between, on the one hand, an almost wild-type number of HEI10/MLH1 foci and the formation of at least one focus per chromosome, and on the other hand the presence of numerous univalents and a large reduction in COs. Further, increasing the numbers of HEI10/MLH1 foci in *heip1* did not increase COs. This supports the idea that AtHEIP1 has an important function in CO maturation, downstream of MLH1 foci formation. In rice *heip1*, the number of late, large HEI10 foci and COs are drastically reduced. Similarly, in mice *c12orf40/Redic1*, the loss of HEIP1 drastically reduces MLH1 foci formation. This suggests that CO designation is more affected by the absence of HEIP1 in these species than in Arabidopsis (30, 31, 40). We thus propose that HEI1P has a pervasive and conserved role in CO formation, with functions both in the designation and maturation of COs. The relative importance of these two functions may vary from species to species.

## Materials and Methods

### Genetic material

The following Arabidopsis lines were used in this study: *heip1-1* (N532319), *heip1-2* (N626287), *heip1-3* (N416767), *msh5-2* (N526553)(45), *hei10-2* (N514624) (26), *mlh1-2* (N1008089) (26), *mus81-2* (N607515) (17), *fancm-1 (33), HEI10-OE* C2 line (*HEI10-OE*) (29), and *zyp1-1(15*). The *heip1-4* and *heip1-5* two full deletion lines were generated by the CRISPR-cas9 system using six guide RNAs targeting the 5’ and 3’ untranslated regions of the *Arabidopsis HEIP1 gene* in Col.0 and *Ler* ecotypes, respectively (Sup dataset 1).

### Cytological techniques

Meiotic chromosome spreads on anthers were performed as previously described (46). Chromosomes were stained with DAPI (1 μg/ml). Images were acquired and processed using a ZEISS microscope (AXIO-Imager.Z2) under a 100× oil immersion objective with ZEN software, and figures were prepared using Adobe Photoshop. Immunolocalization was performed on cells with preserved three-dimensional structures as described in (47). The primary antibodies used were used for both epifluorescence and super-resolution microscopy as follows: anti-REC8 raised in rat (laboratory code PAK036, dilution 1:250), anti-MLH1 raised in rabbit (PAK017, 1:400), and anti-HEI10 raised in chicken (PAK046, 1:50,000). Secondary antibodies were conjugated with Abberior StarRed (STRED-1007-500UG), STAROrange (STORANGE-1005-500UG) and Abberior Stargreen (STGREEN-1002-500UG) for StarRed and STAROrange for STED microscopy (figure 4 and 6) and a Leica THUNDER Imager microscope (Figure 5). MLH1–HEI10 co-foci images were taken with a Leica THUNDER Imager microscope and deconvolved and analyzed with the Imaris software. Super-resolution images were acquired with the Abberior instrument facility line (https://abberior-instruments.com/) using 561- and 640-nm excitation lasers (for STAROrange and STAR Red, respectively) and a 775-nm STED depletion laser. Confocal images were taken with the same instrument with a 485-nm excitation laser (for Alexa 488). Images were deconvolved with Huygens Essential version 20.04 (Scientific Volume Imaging, https://svi.nl/) using the classic maximum likelihood estimation algorithm with lateral drift stabilization; signal-to-noise ratio: 7 for STED images and 20 for confocal images, 40 iterations, and quality threshold of 0.5. Maximum intensity projections and contrast adjustments were done with Huygens Essential.

### CO and aneuploidy identification and analysis by whole-genome sequencing

Plants heterozygous for the *heip1-4* mutation (Col.0) were crossed as females with plants heterozygous for *the heip1-5* mutation (*Ler*). Wild type and *heip1-4/heip1-5* plants were selected among the F1s and crossed as male or as female with wild-type Col.0. Leaf samples from the four back-crossed populations were used for DNA purification and library preparation (48) for Illumina sequencing (HiSeq 3000 2 × 150 base pairs [bp]), performed at the Max Planck-Genome-center (https://mpgc.mpipz.mpg.de/home/).

For the female and male populations, a total of 47 and 48 wild type, 314 and 246 *heip1* plants were sequenced, respectively. The generation of high-confidence SNP markers between Col and *Ler*, mapping of sequencing reads, meiotic CO prediction, filtering of the poorly covered and potentially contaminated samples, and aneuploidy detection were performed as previously described (28). Identified COs were manually and randomly checked by using inGAP-family(49, 50). A total of nine samples were filtered out, two because of a failure in sequencing, three of them were sequenced but with a very low number of reads, three showed signs of inter-sample contamination, while one was derived from selfing according to the genotype. The number of COs per transmitted chromatid in female and male gametes of wild type fom this study were similar from the results from a previous study (49) (Figure S4). The wild-type data from both studies were combined for further analyses of CO number, distribution and interference, as described previously (28).

### CRISPR mutagenesis

The TEFOR website (http://crispor.tefor.net) was used to design guide RNAs that specifically targeted the *HEIP* genes. Six guide RNAs were synthesized with individual U6 promoters in a single cassette (sup data 1) flanked by gateway recombination sites. All guides were cloned into the pDE–Cas9–DSred vector (51, 52) and transformed into wild-type Arabidopsis plants by floral dipping (53). Transgenic plants (T1) were selected based on the seed coat RFP fluorescence marker. To boost the effectiveness of mutagenesis, young plantlets were subjected to heat cycles (54). Then, selected T1 plants were screened for *heip1* deletions by PCR. Subsequently, T2 seeds devoid of fluorescence were chosen as cas9-free and again genotyped for deletion at the *heip1* locus. Deletion at the *HEIP1* locus was confirmed by PCR Sanger sequencing.

## SUPPLEMENTARY DATA & Figure legends

**Figure S1. Metaphase I chromosome spreads of female meiocytes.**

(A) Female wild-type meiocyte with five bivalents. (B) Female *heip1-1* meiocyte with three bivalents. Scale bar, 10 μm. (C) Quantification of bivalents at female metaphase I. Each dot represents the bivalent number of an individual meiotic cell. The mean bivalent number for each genotype is shown by a black bar.

**Figure S2. Colocalization of MLH1 and HEI10 foci in *heip1 HEI10-OE*.**

Immunolocalization of REC8, HEI10, and MLH1 on male meiocytes at late pachytene in *heip1-1 HEI10-OE*. (A) REC8 protein decorates the homologous axis, which are synapsed all along their length. (B) HEI10 foci (C). Merge of REC8 and HEI10 signals. HEI10 foci located in between the two axes. (D) MLH1 foci. (E) HEI10 and MLH1 foci colocalize. Scale bar = 1 μm.

**Figure S3. Analysis of ploidy with sequencing depth and allele ratio along chromosomes.**

Representative samples are shown. The sequencing depth (A, C, E, G, I, M, O), and Col/Ler allelic ratio (B, D, F, H, J) were calculated for each 50 kb interval (with 25 kb step size) along chromosomes. The pericentromeric and centromeric regions are indicated by grey and blue shading, respectively. On the top panel, the horizontal dashed line indicates the mean sequencing depth of the sample. On the bottom panel, red and blue dots correspond to Col and Ler allelic frequencies, respectively.

(A,B) A sample derived from wild-type female hybrid (5385_AW). The coverage is even along and among chromosomes, suggesting euploidy. The allelic ratio suggests regions of heterozygosity (similar frequency of Col.0 and Ler polymorphisms) and Col homozygositysity, the transitions corresponding to crossovers.

(C,D) A sample derived from wild-type female hybrid (5385_AV). The even coverageand allelic ratios suggest euploidy.

(E,F) A sample derived from *heip1* female hybrid (5388_BT). The even coverage and allelic ratios suggest euploidy.

(G, H) A sample derived from *heip1* male hybrid (5388_BU). The even coverage and allelic ratios suggest euploidy.

(I, J) A sample derived from *heip1* female hybrid (5388_D). Chromosome 4 has ~50% more coverage than the other chromosomes, suggesting trisomy 4. The Col/Ler allelic ratio of ~2:1 along chromosome 4 also supports this conclusion.

(K, L). A sample derived from *heip1* male hybrid (5389_R). Chromosome 2 has ~50% more coverage than the other chromosomes, suggesting trisomy 2. The Col/Ler allelic ratio of ~2:1 along chromosome 4 also supports this conclusion.

(M, N) A sample derived from *heip1* female hybrid (5388_AT). The left arm of Chromosome 5 has ~50% more coverage than the right arm and the other chromosomes, suggesting segmental trisomy. The Col/Ler allelic ratio of ~2:1 along the left arm of chromosome 5 also supports this conclusion.

(O, P) A sample derived from *heip1* female hybrid. Two segments of chromosome 4 have higher coverage and a 2:1 allelic ratio, suggesting a complex genome rearrangement.rearrangement

**Figure S4. Analysis of CO number in wild type**

The number of COs per transmitted chromatid in female and male gametes of wild type from a previous study (49) versus this study (sister plants of *heip1* mutants are not different). Fisher’s LSD test was applied. The wild-type data from a previous study was further analysed in this study.

## Acknowledgments

We would like to thank the Max Planck Genome center for library preparation and sequencing. This work was supported by core funding from the Max Planck Society and an Alexander von Humboldt Fellowship to Q.L. This work has benefited from the support of IJPB’s Plant Observatory technological platforms. The IJPB benefits from the support of Saclay Plant Sciences-SPS (ANR-17-EUR-0007). We thank Neysan Donnelly for proofreading the manuscript.

## Notes

### Competing Interest Statement

The authors have declared no competing interest.

## Bibliography

1. T. Hassold, H. Hall, P. Hunt, The origin of human aneuploidy: where we have been, where we are going. Hum Mol Genet 16 Spec No. 2, R203–208 (2007).

2. M. Herbert, D. Kalleas, D. Cooney, M. Lamb, L. Lister, Meiosis and maternal aging: insights from aneuploid oocytes and trisomy births. Cold Spring Harb Perspect Biol 7, a017970 (2015).

3. N. Hunter, Meiotic Recombination: The Essence of Heredity. Cold Spring Harb Perspect Biol 7 (2015).

4. B. de Massy, Initiation of meiotic recombination: how and where? Conservation and specificities among eukaryotes. Annu Rev Genet 47, 563–599 (2013).

5. Y. Wang, G. P. Copenhaver, Meiotic Recombination: Mixing It Up in Plants. Annu Rev Plant Biol 69, 577–609 (2018).

6. S. Gray, P. E. Cohen, Control of Meiotic Crossovers: From Double-Strand Break Formation to Designation. Annu Rev Genet 50, 175–210 (2016).

7. J. L. Youds, S. J. Boulton, The choice in meiosis - defining the factors that influence crossover or non-crossover formation. J Cell Sci 124, 501–513 (2011).

8. T. de los Santos et al., The Mus81/Mms4 endonuclease acts independently of double-Holliday junction resolution to promote a distinct subset of crossovers during meiosis in budding yeast. Genetics 164, 81–94 (2003).

9. S. Wang, D. Zickler, N. Kleckner, L. Zhang, Meiotic crossover patterns: obligatory crossover, interference and homeostasis in a single process. Cell Cycle 14, 305–314 (2015).

10. L. von Diezmann, O. Rog, Let’s get physical - mechanisms of crossover interference. J Cell Sci 134 (2021).

11. K. Zakharyevich, S. Tang, Y. Ma, N. Hunter, Delineation of joint molecule resolution pathways in meiosis identifies a crossover-specific resolvase. Cell 149, 334–347 (2012).

12. N. Jackson et al., Reduced meiotic crossovers and delayed prophase I progression in AtMLH3-deficient Arabidopsis. EMBO J 25, 1315–1323 (2006).

13. E. Cannavo et al., Regulation of the MLH1-MLH3 endonuclease in meiosis. Nature 586, 618–622 (2020).

14. L. Ranjha, R. Anand, P. Cejka, The Saccharomyces cerevisiae Mlh1-Mlh3 heterodimer is an endonuclease that preferentially binds to Holliday junctions. J Biol Chem 289, 5674–5686 (2014).

15. L. Capilla-Perez et al., The synaptonemal complex imposes crossover interference and heterochiasmy in Arabidopsis. Proc Natl Acad Sci U S A 118 (2021).

16. L. Chelysheva et al., An easy protocol for studying chromatin and recombination protein dynamics during Arabidopsis thaliana meiosis: immunodetection of cohesins, histones and MLH1. Cytogenet Genome Res 129, 143–153 (2010).

17. L. E. Berchowitz, K. E. Francis, A. L. Bey, G. P. Copenhaver, The role of AtMUS81 in interference-insensitive crossovers in A. thaliana. PLoS Genet 3, e132 (2007).

18. L. J. Gaskell, F. Osman, R. J. Gilbert, M. C. Whitby, Mus81 cleavage of Holliday junctions: a failsafe for processing meiotic recombination intermediates? EMBO J 26, 1891–1901 (2007).

19. G. V. Borner, N. Kleckner, N. Hunter, Crossover/noncrossover differentiation, synaptonemal complex formation, and regulatory surveillance at the leptotene/zygotene transition of meiosis. Cell 117, 29–45 (2004).

20. Q. Zhang, S. Y. Ji, K. Busayavalasa, C. Yu, SPO16 binds SHOC1 to promote homologous recombination and crossing-over in meiotic prophase I. Sci Adv 5, eaau9780 (2019).

21. A. De Muyt et al., A meiotic XPF-ERCC1-like complex recognizes joint molecule recombination intermediates to promote crossover formation. Genes Dev 32, 283–296 (2018).

22. R. Mercier, C. Mezard, E. Jenczewski, N. Macaisne, M. Grelon, The molecular biology of meiosis in plants. Annu Rev Plant Biol 66, 297–327 (2015).

23. D. Zickler, N. Kleckner, Recombination, Pairing, and Synapsis of Homologs during Meiosis. Cold Spring Harb Perspect Biol 7 (2015).

24. M. G. France et al., ZYP1 is required for obligate cross-over formation and cross-over interference in Arabidopsis. Proc Natl Acad Sci U S A 118 (2021).

25. K. Wang et al., The role of rice HEI10 in the formation of meiotic crossovers. PLoS Genet 8, e1002809 (2012).

26. L. Chelysheva et al., The Arabidopsis HEI10 Is a New ZMM Protein Related to Zip3. PLOS Genetics 8, e1002799 (2012).

27. C. Morgan et al., Diffusion-mediated HEI10 coarsening can explain meiotic crossover positioning in Arabidopsis. Nat Commun 12, 4674 (2021).

28. S. Durand et al., Joint control of meiotic crossover patterning by the synaptonemal complex and HEI10 dosage. Nat Commun 13, 5999 (2022).

29. P. A. Ziolkowski et al., Natural variation and dosage of the HEI10 meiotic E3 ligase control Arabidopsis crossover recombination. Genes Dev 31, 306–317 (2017).

30. Y. Li et al., HEIP1 regulates crossover formation during meiosis in rice. Proc Natl Acad Sci U S A 115, 10810–10815 (2018).

31. Z. Chang et al., The plant-specific ABERRANT GAMETOGENESIS 1 gene is essential for meiosis in rice. J Exp Bot 71, 204–218 (2020).

32. J. D. Higgins, E. F. Buckling, F. C. Franklin, G. H. Jones, Expression and functional analysis of AtMUS81 in Arabidopsis meiosis reveals a role in the second pathway of crossing-over. Plant J 54, 152–162 (2008).

33. W. Crismani et al., FANCM limits meiotic crossovers. Science 336, 1588–1590 (2012).

34. W. Guo, L. Comai, I. M. Henry, Chromoanagenesis in the asy1 meiotic mutant of Arabidopsis. G3 (Bethesda) 10.1093/g3journal/jkac185 (2022).

35. S. Wang et al., Inefficient Crossover Maturation Underlies Elevated Aneuploidy in Human Female Meiosis. Cell 168, 977–989 e917 (2017).

36. L. Zimmermann et al., A Completely Reimplemented MPI Bioinformatics Toolkit with a New HHpred Server at its Core. J Mol Biol 430, 2237–2243 (2018).

37. F. Gabler et al., Protein Sequence Analysis Using the MPI Bioinformatics Toolkit. Curr Protoc Bioinformatics 72, e108 (2020).

38. J. Jumper et al., Highly accurate protein structure prediction with AlphaFold. Nature 596, 583–589 (2021).

39. M. Varadi et al., AlphaFold Protein Structure Database: massively expanding the structural coverage of protein-sequence space with high-accuracy models. Nucleic Acids Res 50, D439–D444 (2022).

40. Q. Shi et al., A novel recombination protein C12ORF40/REDIC1 is required for meiotic crossover formation. Research Square 10.21203/rs.3.rs-1971629/v1 (2022).

41. N. Macaisne, J. Vignard, R. Mercier, SHOC1 and PTD form an XPF-ERCC1-like complex that is required for formation of class I crossovers. J Cell Sci 124, 2687–2691 (2011).

42. P. E. Wright, H. J. Dyson, Intrinsically disordered proteins in cellular signalling and regulation. Nat Rev Mol Cell Biol 16, 18–29 (2015).

43. D. C. Nageswaran et al., HIGH CROSSOVER RATE1 encodes PROTEIN PHOSPHATASE X1 and restricts meiotic crossovers in Arabidopsis. Nat Plants 7, 452–467 (2021).

44. J. Loidl, Conservation and Variability of Meiosis Across the Eukaryotes. Annu Rev Genet 50, 293–316 (2016).

45. J. D. Higgins et al., AtMSH5 partners AtMSH4 in the class I meiotic crossover pathway in Arabidopsis thaliana, but is not required for synapsis. Plant J 55, 28–39 (2008).

46. K. J. Ross, P. Fransz, G. H. Jones, A light microscopic atlas of meiosis in Arabidopsis thaliana. Chromosome Res 4, 507–516 (1996).

47. A. Hurel et al., A cytological approach to studying meiotic recombination and chromosome dynamics in Arabidopsis thaliana male meiocytes in three dimensions. Plant J 95, 385–396 (2018).

48. B. A. Rowan, V. Patel, D. Weigel, K. Schneeberger, Rapid and inexpensive whole-genome genotyping-by-sequencing for crossover localization and fine-scale genetic mapping. G3 (Bethesda) 5, 385–398 (2015).

49. Q. Lian et al., The megabase-scale crossover landscape is largely independent of sequence divergence. Nat Commun 13, 3828 (2022).

50. Q. Lian, Y. Chen, F. Chang, Y. Fu, J. Qi, inGAP-family: Accurate Detection of Meiotic Recombination Loci and Causal Mutations by Filtering Out Artificial Variants due to Genome Complexities. Genomics Proteomics Bioinformatics 10.1016/j.gpb.2019.11.014 (2021).

51. F. Fauser, S. Schiml, H. Puchta, Both CRISPR/Cas-based nucleases and nickases can be used efficiently for genome engineering in Arabidopsis thaliana. Plant J 79, 348–359 (2014).

52. C. Morineau et al., Selective gene dosage by CRISPR-Cas9 genome editing in hexaploid Camelina sativa. Plant Biotechnol J 15, 729–739 (2017).

53. X. Zhang, R. Henriques, S. S. Lin, Q. W. Niu, N. H. Chua, Agrobacterium-mediated transformation of Arabidopsis thaliana using the floral dip method. Nat Protoc 1, 641–646 (2006).

54. C. LeBlanc et al., Increased efficiency of targeted mutagenesis by CRISPR/Cas9 in plants using heat stress. Plant J 93, 377–386 (2018).

