## Supplementary material for "HEIP1 is required for efficient meiotic crossover implementation and is conserved from plants to humans": Figure S1-S4

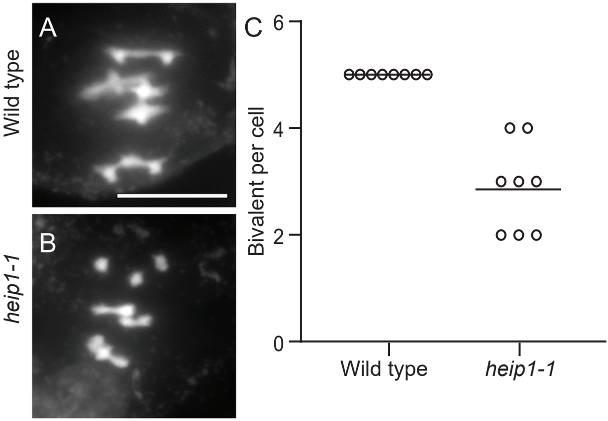


Figure S1.

Metaphase I chromosome spreads of female meiocytes. (A) Female wild-type meiocyte with five bivalents. (B) Female heip1-1 meiocyte with three bivalents. Scale bar, 10 µm. (C) Quantification of bivalents at female metaphase I. Each dot represents the bivalent number of an individual meiotic cell. The mean bivalent number for each genotype is shown by a black bar.

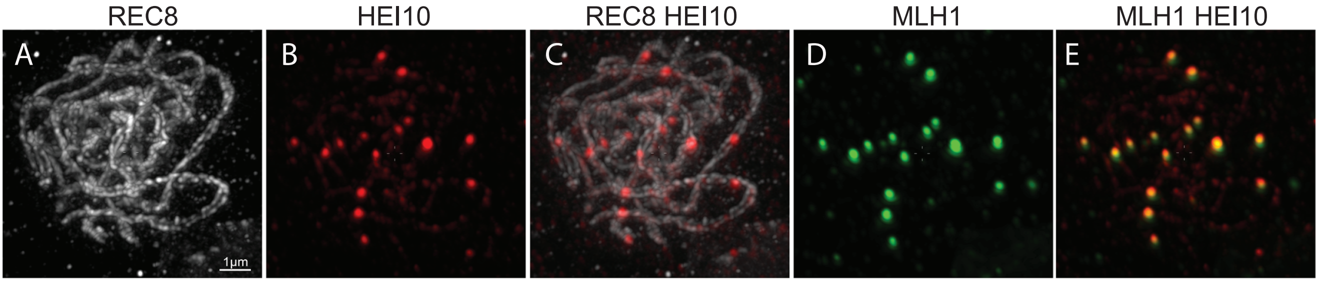


Figure S2.

Colocalization of MLH1 and HEI10 foci in heip1 HEI10-OE.
Immunolocalization of REC8, HEI10, and MLH1 on male meiocytes at late pachytene in heip1-1 HEI10-OE. (A) REC8 protein decorates the homologous axis, which are synapsed all along their length. (B) HEI10 foci (C). Merge of REC8 and HEI10 signals. HEI10 foci located in between the two axes. (D) MLH1 foci. (E) HEI10 and MLH1 foci colocalize. Scale bar = 1 µm.

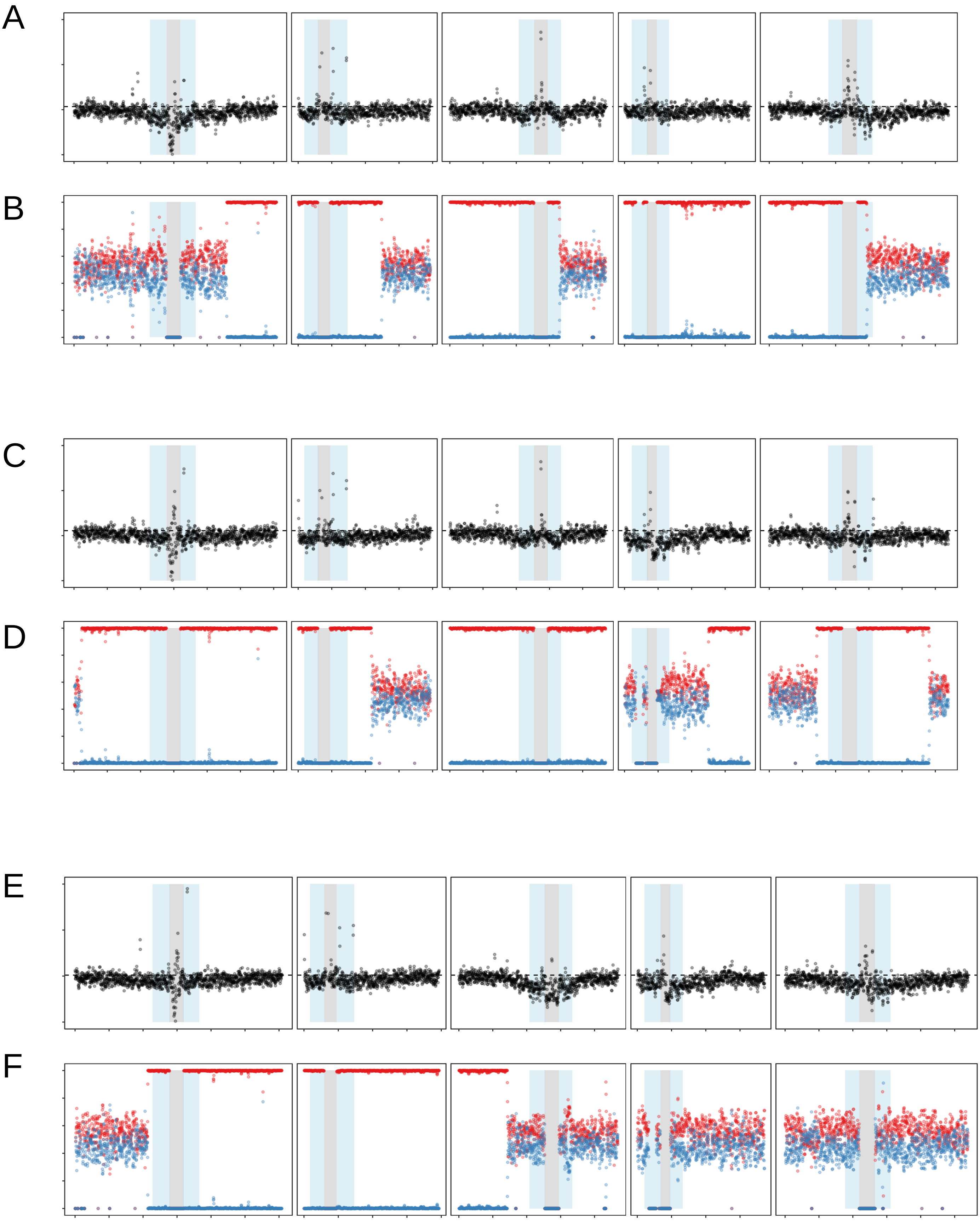
A
B
C
D
E
F

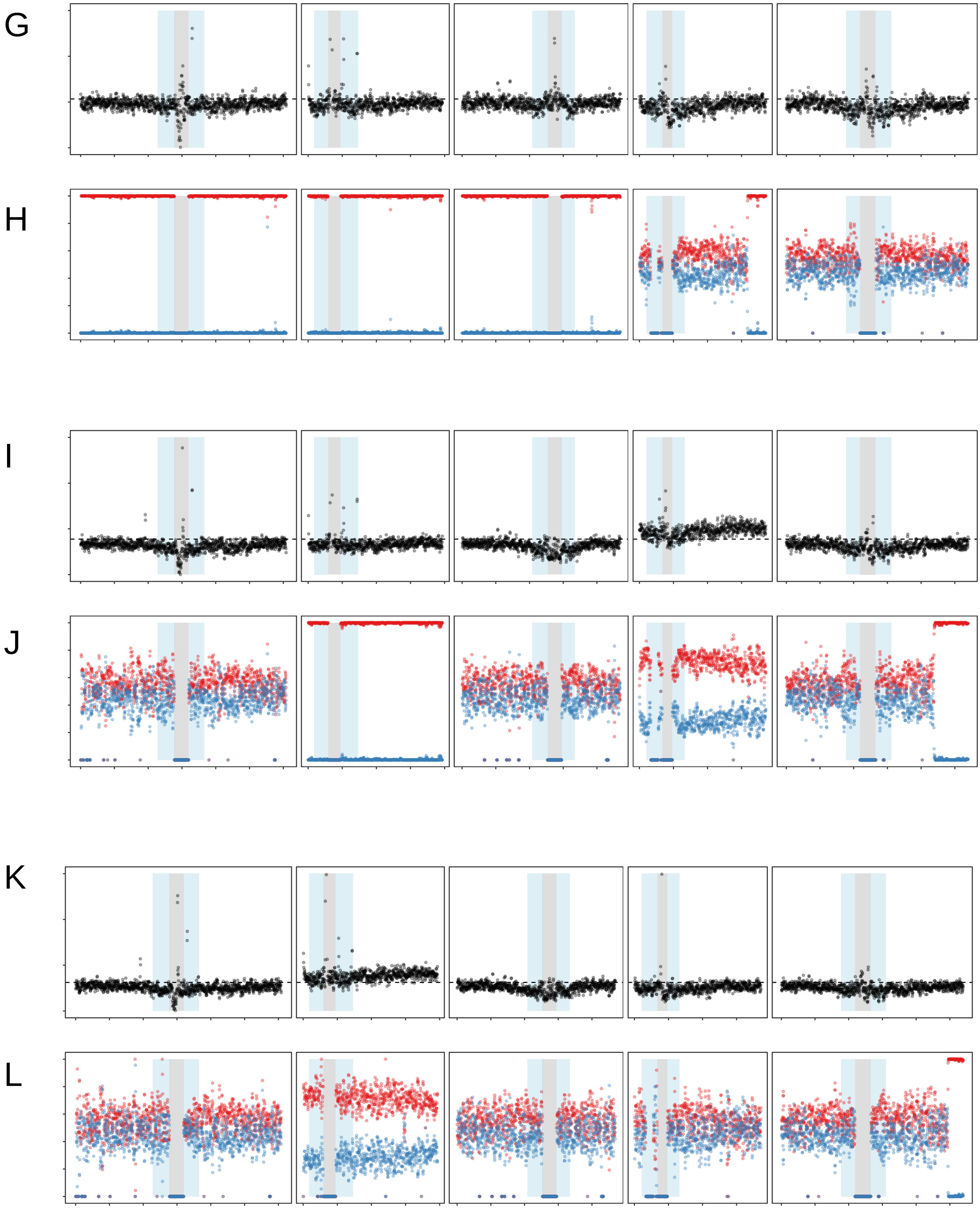
G
H
I
J
K
L

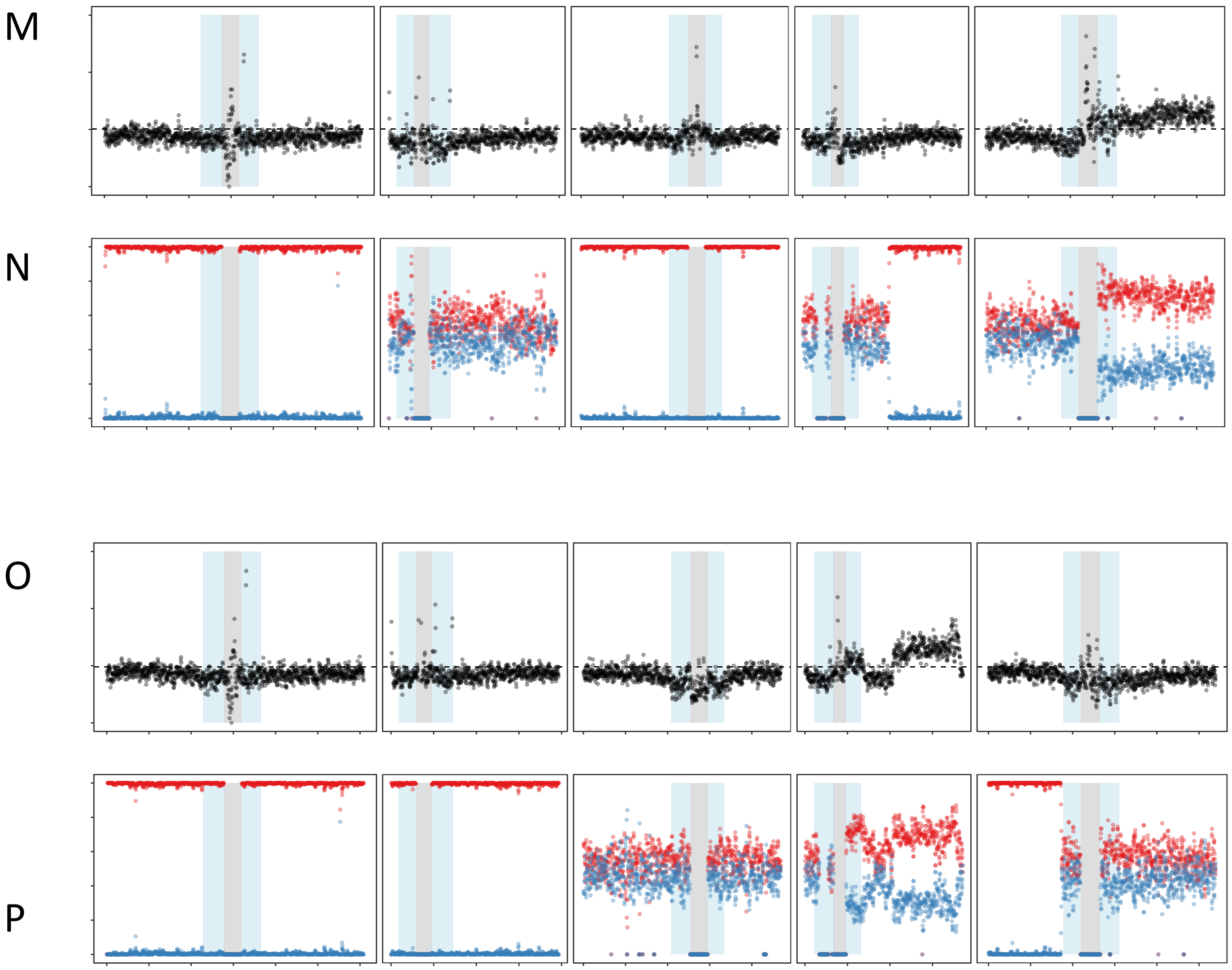


**Figure S3..**

Analysis of ploidy with sequencing depth and allele ratio along chromosomes.
Representative samples are shown. The sequencing depth (A, C, E, G, I, M, O), and Col/Ler allelic ratio (B, D, F, H, J) were calculated for each 50 kb interval (with 25 kb step size) along chromosomes. The pericentromeric and centromeric regions are indicated by grey and blue shading, respectively. On the top panel, the horizontal dashed line indicates the mean sequencing depth of the sample. On the bottom panel, red and blue dots correspond to Col and Ler allelic frequencies, respectively.

(A,B) A sample derived from wild-type female hybrid (5385_AW). The coverage is even along and among chromosomes, suggesting euploidy. The allelic ratio suggests regions of heterozygosity (similar frequency of Col.0 and Ler polymorphisms) and Col homozygositysity, the transitions corresponding to crossovers.
(C,D) A sample derived from wild-type female hybrid (5385_AV). The even coverageand allelic ratios suggest euploidy.
(E,F) A sample derived from heip1 female hybrid (5388_BT). The even coverage and allelic ratios suggest euploidy.
(G, H) A sample derived from heip1 male hybrid (5388_BU). The even coverage  and allelic ratios suggest euploidy.
(I, J) A sample derived from heip1 female hybrid (5388_D). Chromosome 4 has ~50% more coverage than the other chromosomes, suggesting trisomy 4. The Col/Ler allelic ratio of ~2:1 along chromosome 4 also supports this conclusion.
(K, L). A sample derived from heip1 male hybrid (5389_R). Chromosome 2 has ~50% more coverage than the other chromosomes, suggesting trisomy 2. The Col/Ler allelic ratio of ~2:1 along chromosome 4 also supports this conclusion.
(M, N) A sample derived from heip1 female hybrid (5388_AT). The left arm of Chromosome 5 has ~50% more coverage than the right arm and the other chromosomes, suggesting segmental trisomy. The Col/Ler allelic ratio of ~2:1 along the left arm of chromosome 5 also supports this conclusion.
(O, P) A sample derived from heip1 female hybrid. Two segments of chromosome 4 have higher coverage and a 2:1 allelic ratio, suggesting a complex genome rearrangement.rearrangement

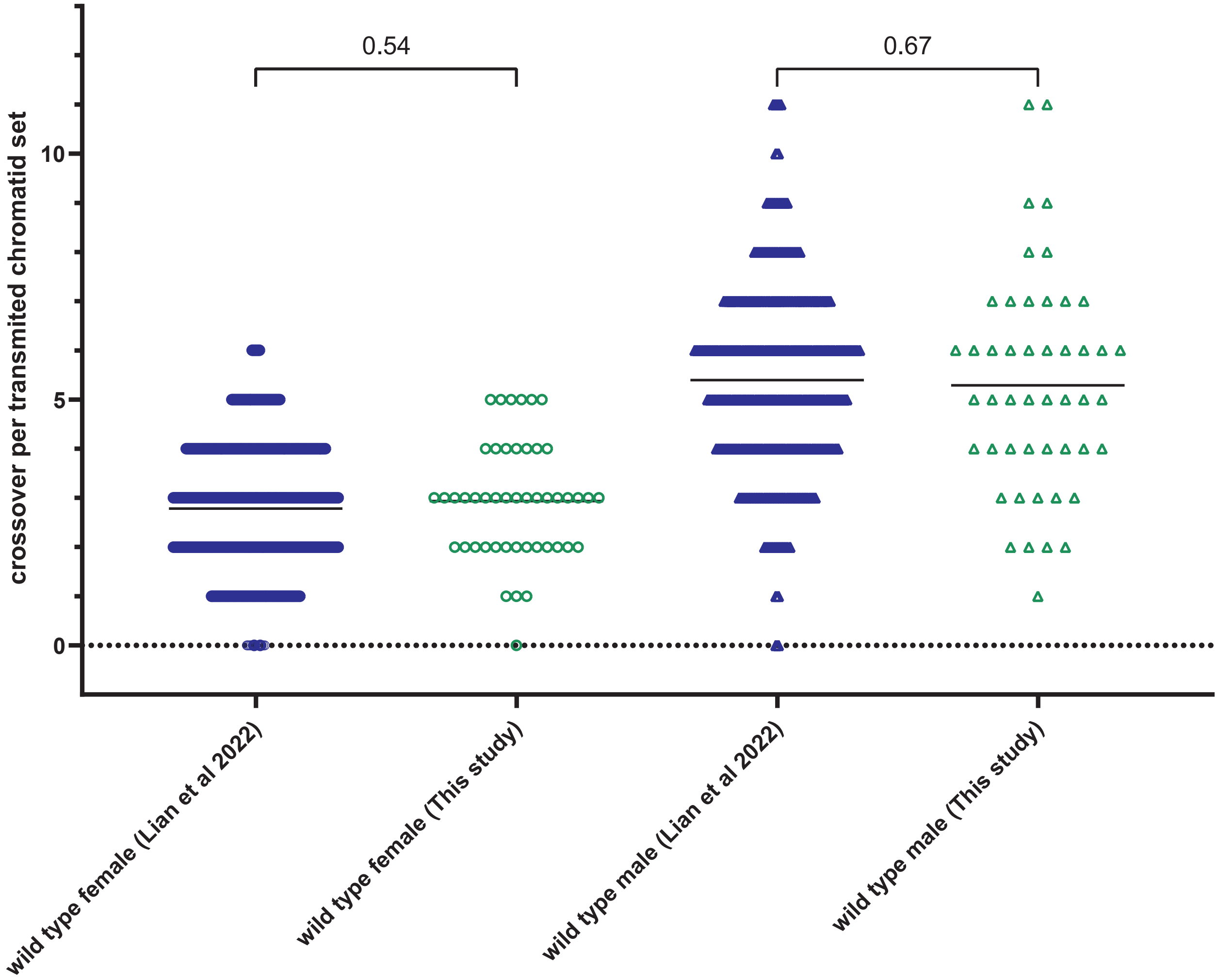


Figure S4

Analysis of CO number in wild type
The number of COs per transmitted chromatid in female and male gametes of wild type from a previous study (48) versus this study (sister plants of heip1 mutants are not different). Fisher's LSD test was applied. The wild-type data from a previous study was further analysed in this study.
